# Automatic Cell Segmentation by Adaptive Thresholding (ACSAT) for large scale calcium imaging datasets

**DOI:** 10.1101/260075

**Authors:** Simon P. Shen, Hua-an Tseng, Kyle R. Hansen, Ruofan Wu, Howard Gritton, Jennie Si, Xue Han

**Affiliations:** Department of Physics, Harvard University, Cambridge, MA 02138; Biomedical Engineering Department, Boston University, Boston, MA 02215; School of Electrical, Computer and Energy Engineering, Tempe, AZ 85287

## Abstract

Advances in calcium imaging have made it possible to record from an increasingly larger number of neurons simultaneously. Neuroscientists can now routinely image hundreds to thousands of individual neurons. With the continued neurotechnology development effort, it is expected that millions of neurons could soon be simultaneously measured. An emerging technical challenge that parallels the advancement in imaging such a large number of individual neurons is the processing of correspondingly large datasets, an important step of which is the identification of individual neurons. Traditional methods rely mainly on manual or semi-manual inspection, which cannot be scaled to processing large datasets. To address this challenge, we have developed an automated cell segmentation method, which is referred to as Automated Cell Segmentation by Adaptive Thresholding (ACSAT). ACSAT includes an iterative procedure that automatically calculates global and local threshold values during each iteration based on image pixel intensities. As such, the algorithm is capable of handling morphological variations and dynamic changes in fluorescence intensities in different calcium imaging datasets. In addition, ACSAT computes adaptive threshold values based on a time-collapsed image that is representative of the image sequence, and thus ACSAT provides segmentation results at a fast speed. We tested the algorithm on wide-field calcium imaging datasets in the brain regions of hippocampus and striatum in mice. ACSAT achieved precision and recall rates of approximately 80% when compared to individual neurons that are verified by human inspection. Additionally, ACSAT successfully detected low-intensity neurons that were initially undetected by humans.

**Significance:** ACSAT automatically segments cells in large scale wide-field calcium imaging datasets. It is based on adaptive thresholding at both global and local levels, implemented in an iterative process to identify individual neurons in a time-collapsed image from an image sequence. It is therefore capable of handling variation in cell morphology and dynamic changes between different calcium imaging datasets at a fast speed. Based on tests performed on two datasets from mouse hippocampus and striatum, ACSAT performed comparable to human referees and was even able to detect low-intensity neurons that were initially undetected by human referees.

## Introduction

The ability to record from a large population of single neurons during behavior greatly facilitates the investigation of the contribution of individual neurons to neural network dynamics. The extracellular single-unit recording technique has traditionally been a method of choice in neurophysiological analysis of single neurons in the brain. Recent improvements in single-cell imaging techniques using activity indicators have offered new alternatives. Voltage sensors (Ghitani et al., 2015, Gong et al., 2015, Chamberland et al., 2017, Inagaki et al., 2017) and calcium sensors (Chen et al., 2013, Sun et al., 2013) have made it possible to observe hundreds to thousands of individual neurons simultaneously (Ohki et al., 2005, Andermann et al., 2010, Huber et al., 2012, Ziv et al., 2013, Issa et al., 2014, Mohammed et al., 2016). Though indirect, calcium indicators, especially the newest generation of the genetically encoded calcium sensor GCaMP6 (Chen et al., 2013), have been sensitive enough to monitor neural activity with high spatiotemporal precision in behaving animals, allowing researchers to examine the activity of populations of a specific cell type (Hofer et al., 2011, Wachowiak et al., 2013, Pinto and Dan, 2015, Allen et al., 2017), or the same cell over an extended period of time (Poort et al., 2015).

As the signal-to-noise ratio of the genetically encoded calcium indicators improved, wide-field microscopy has become a feasible choice for recording the activity of a large population of neurons over an extended anatomical area. Although lacking the spatial subcellular resolution of a multiphoton microscope, wide-field microscopes can operate at a higher speed and allow the simultaneous recording of a larger population, (Ghosh et al., 2011, Ziv et al., 2013, Kim et al., 2016, Mohammed et al., 2016). Advanced microfabrication techniques further miniaturized the wide-field microscope to an microendoscope, capable of monitoring neural activity in freely-moving animals (Ghosh et al., 2011, Ziv et al., 2013).

An emerging technical challenge that parallels advances in imaging of increasingly more neurons is the processing of large datasets. During data analysis, an important step is to identify regions of interest (ROIs) corresponding to individual neurons. As data grows rapidly both spatially and temporally, the traditional labor-intensive approach of manual inspection has to be replaced by semi- or fully-automatic procedures. Principal component analysis (PCA) (Mukamel et al., 2009), as one such approach, requires significant computational resources and CPU processing time, limiting its use in larger datasets.

Alternatively, threshold-based methods are simple, intuitive, and fast, and thus are expected to be useful for processing large datasets. However, before applying threshold-based methods to calcium-imaging datasets, several challenges need to be overcome, including: inhomogeneity across the imaging field, variations in recording condition, recording subjects, and fluorescence signal strength. For example, one of the most referenced thresholding methods, Otsu’s method, which automatically selects the optimal threshold value that minimizes the intra-class variance among ROI pixels and among background pixels, would only successfully segment some of the highest-intensity ROIs (Otsu, 1979, Sezgin and Sankur, 2004). Additionally, the multi-class Otsu’s method is limited when identifying ROIs with different pixel intensities due to the uneven lighting of the background. A recent machine learning-based algorithm uses image gradients and pixel traces to automatically optimize the threshold value (Fantuzzo et al., 2017). However, the method still requires a user’s subjective input in selecting a background removal factor based on the dataset. Other approaches based on edge detection have trouble due to weak fluorescence signal strength in comparison with the background pixels (Sadeghian et al., 2009). Even if edges were detected, it remains difficult to objectively determine which edges belong to which ROIs because of the high density of neurons in the image. In general, most segmentation methods require a high level of tuning to each individual dataset and individualized calibration to establish thresholds, e.g. for gradient values or for pixel intensity values. Thus, thresholding methods have shown promise for automatic cell segmentation if the above discussed technical challenges can be addressed to overcome variability among and within datasets.

We introduce a new Automated Cell Segmentation by Adaptive Thresholding (ACSAT) algorithm, which dynamically and automatically determines global and local threshold values based on pixel intensity levels within a time-collapsed image of a recorded image sequence. We applied ACSAT to two datasets collected from mice hippocampus and striatum, which have distinct cell morphology, calcium dynamics, and fluorescence intensity levels. ACSAT achieved precision and recall rates of approximately 80% when compared to ROIs that can be verified by human referees manually, and was successful at identifying low-intensity neurons that were initially undetected by human referees.

## Materials and Methods

### Mouse preparation

All animal procedures were approved by Boston University Institutional Animal Care and Use Committee. Female C57BL/6 mice (8-12 weeks old, Taconic, Hudson, NY) were first injected with 250nL AAV9-Syn-GCaMP6.WPRE.SV40 virus (titer: ~6e12 GC/ml, University of Pennsylvania Vector Core). AAV was delivered either into the dorsal CA1 (AP: -2, ML: 1.4, DV: -1.6), or into the dorsal striatum (AP: 0.5, ML: 1.8, DV: -1.6) regions. Injections were performed with a 10 μL syringe (World Precision Instruments, Sarasota, FL) coupled with a 33 gauge needle (NF33BL, World Precision Instruments, Sarasota, FL) at a speed of 40 nL/min, controlled by a microsyringe pump (UltraMicroPump 3-4, World Precision Instruments, Sarasota, FL). Upon complete recovery, a custom imaging chamber with glass coverslip was surgically implanted on top of the viral injection site by removing the overlying cortical tissue. The imaging chamber was assembled by fitting a circular coverslip (size 0; OD: 3 mm) to a stainless steel cannula (OD: 0.317 mm, ID: 0.236 mm) using a UV-curable optical adhesive (Norland Products). During surgery, a custom aluminum headplate was also attached to the skull, which allows head-fixation during the imaging session.

The hippocampus dataset was previously reported by Mohammed et al. (2016). In this dataset, the mouse was trained to perform a trace conditioning task known to involve hippocampal neural activity (Solomon et al., 1986, Moyer et al., 1990, Tseng et al., 2004, Sakamoto et al., 2005). In this task, the animal was trained to associate a conditioned stimulus (a 350 ms long tone) with an unconditioned stimulus (a gentle 100ms air puff to one eye). There was a 250 ms trace interval between two stimuli. During each recording session, the animal was head-fixed and performed 40 trials with a randomized 31-36 second inter-trial interval. The hippocampus dataset (1024 × 1024 pixels/frame, 2047 frames, ~4 GB size) analyzed in this study was part of a larger recording session (~ 50GB size).

The striatum dataset was collected from a head-fixed animal running on a spherical treadmill system. The treadmill system consisted a styrofoam ball floated by air pressure in a 3D-printed bowl designed as described in (Dombeck et al., 2007) that allowed the animal to move its limbs freely while head-fixed. The mouse was first handled for several days before being headfixed to the spherical treadmill. Habituation to running on the spherical treadmill while headfixed occurred over 3-4 days/week at the same time of day as subsequent recording sessions (8-12 hours after lights-on), for several weeks. Single imaging sessions took approximately 25 minutes. Sampling occurred at approximately 20Hz and exposure time was fixed at 20ms. The striatum dataset (~4 GB size) contains 2047 frames with 1024 x 1024 pixels per frame and was also part of a larger dataset (~25GB size).

### Calcium imaging data acquisition

Imaging data were acquired with a custom wide-field microscope coupled with a scientific CMOS camera (ORCA-Flash 4.0, C11440-42U, Hamamatsu, Boston MA), controlled by the commercial software package HCImageLive (Hamamatsu, Boston MA). The wide-field microscope consisted of a Leica N Plan 10×0.25 PH1 objective lens, an excitation filter (HQ 470/50), a dichroic mirror (FF506-Di02), an emission filter (FF01-536/40), a commercial SLR lens as the tube lens (Nikon Zoom-NIKKOR 80-200 mm f/4 AI-s), and a 5W LED (LZ1-00B200, 460 nm; LedEngin, San Jose CA). Data acquisition was performed at 20 Hz, at a resolution of 1024 × 1024 pixels, with 16-bits per pixel. With 10x objective lens, the microscope provided a field of view of 1.343 × 1.343 mm^2^ (1.312 × 1.312 μm^2^/pixel) of brain tissue. During data acquisition, head-fixed animals were either performing the trace conditioning task (the hippocampus dataset) or moving on the spherical treadmill (the striatum dataset). Imaging data was streamed from the camera to RAM of a custom computer (dual Intel Xeon processors, 128 GB RAM, and a GeForce GTX Titan video card) to ensure temporal precision. After each imaging session, data was moved from RAM to hard drive and saved in multi-page tagged image file format.

### Automated Cell Segmentation by Adaptive Thresholding (ACSAT) Overview

Fluorescence imaging data obtained in the form of image sequences is processed offline using a custom MATLAB algorithm. Image sequences were first motion-corrected as described in Mohammed et al. (2016) to remove micromotion of the imaged area caused by breathing and other slight movements of the animal. Our Automated Cell Segmentation by Adaptive Thresholding (ACSAT) method (Figure 1a) is then applied to the image sequences. The overall goal is to automatically identify individual neurons as regions of interest (ROIs) from the image sequence so that the activity of each neuron can be approximated by the fluorescence intensity within that ROI.

**Figure 1.**
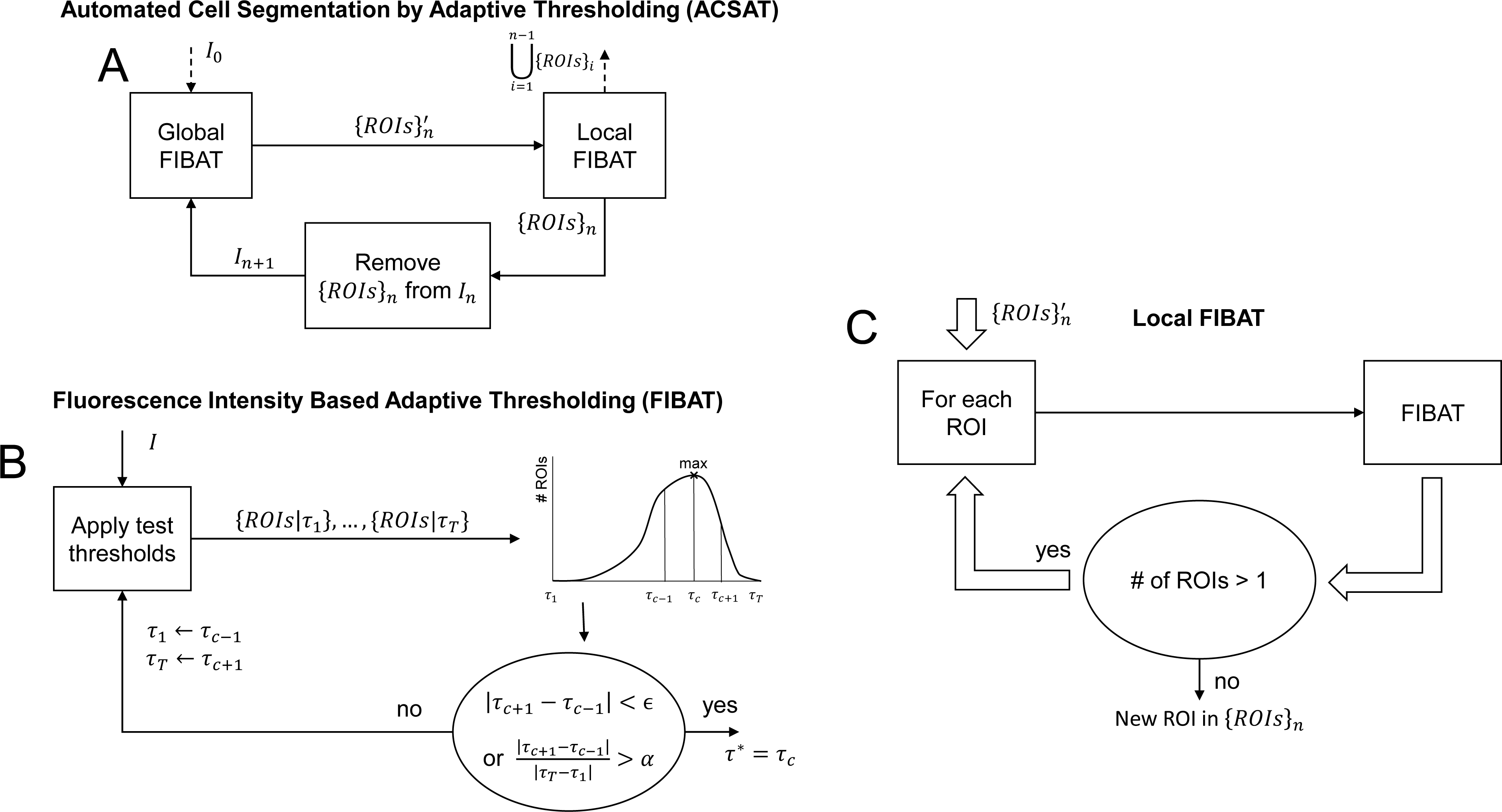
Automated Cell Segmentation by Adaptive Thresholding (ACSAT) algorithm flowchart. (A) The input is the time-collapsed image *I*_0_, and the output is a collection of automatically segmented ROIs. In each iteration, the “Global FIBAT” step identifies potential ROIs 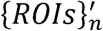 by applying FIBAT, described in part B, to the entire image *I*_*n*_; and the “Local FIBAT” step, described in part C, splits overlapping ROIs. (B) Fluorescence Intensity Based Adaptive Thresholding (FIBAT) algorithm flowchart. The inputted image is segmented using each of the test threshold values *τ*_1_, *τ*_2_,*τ*_3_…, *τ*_*T*_. The search range for the optimal threshold value (*τ*_1_, *τ*_*T*_) is iteratively narrowed to contain the test threshold value which resulted in the maximum number of ROIs. (C) Local FIBAT procedure. FIBAT, described in part B, is recursively applied to each individual inputted ROI until the resulting ROI(s) can no longer be separated by FIBAT.

To increase time efficiency of the ACSAT algorithm without sacrificing segmentation performance, the inputted image sequence is first collapsed in time into one representative two-dimensional image (*I*_0_ in Figure 1a), where each pixel in *I*_0_ is thus represented by the maximum intensity value of that pixel across the entire image sequence with the mean value removed. This image is then used for the rest of the algorithm. Pixels with low intensity values would correspond to static background, whereas pixels with high intensity values would correspond to neurons with GCaMP6 expression. Because we define a ROI as a non-trivial cluster of adjacent pixels with high intensity values in *I*_0_, it is unlikely for a significant cluster of background pixels to all have high intensity values due to random noise. Thus, the time-collapsed image is expected to contain sufficient information to separate neurons from the background.

Next, ACSAT iteratively generates ROIs *{ROIs}*_*n*_ from the time-collapsed image *I*_*n*_ for iterations *n* = 1,2,…, starting with *I*_1_ = *I*_0_. Prior to each subsequent iteration, *I*_*n*_ is generated by cumulatively clearing previously segmented ROIs, *{ROIs}*_*n-1*_, from *I*_*n-l*_ by setting each ROI’s pixels in *I*_*n-1*_ to blank values of 0 and dilating the cleared area. As described later, each iteration consists of both adaptive thresholding at the global level (Global FIBAT in Figure 1a), using the automatically selected threshold value 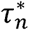 (Figure 1b), and adaptive thresholding at the local level (Local FIBAT in Figure 1a). When the change in global threshold value 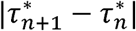 is insignificant, further iterations are likely to contribute more false positives than true positives. Thus, the ACSAT algorithm terminates at iteration *n* if 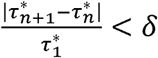 where 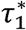 acts as a normalizing factor. In this study, we chose *δ* = 10%. Accordingly, the final output of ACSAT is the union of the segmented ROIs from each iteration, *{ROIs}*_*1*_ ⋃ … ⋃ *{ROIs}*_*n*_.

### Global and Local Adaptive Thresholding in ACSAT

Each iteration *n* of ACSAT contributes a set of newly segmented ROIs *{ROIs}*_*n*_ from *I*_*n*_ by applying our Fluorescence Intensity Based Adaptive Thresholding (FIBAT) algorithm, at the global and local levels (Global/Local FIBAT in Figure 1a). Briefly, FIBAT (Figure 1b) takes an inputted image *I* and outputs the optimal threshold value *τ*\* which results in optimally segmented ROIs *{ROIs*|*τ*\*}.

Global adaptive thresholding is the first step in the nth iteration of ACSAT (Figure 1a). This step applies FIBAT directly to the whole image (*I*_n_ →*I*) to identify potential ROIs 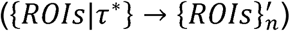.

These potential ROIs 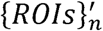 may include groups of adjacent neurons or overlapping neurons simply because neurons physically are located in a 3D space during in vivo imaging. The local adaptive thresholding step (Figure 1c) recursively separates any potentially overlapping ROIs within 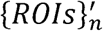 in order to output *{ROIs}*_*n*_. Specifically, each ROI in 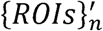 is individually inputted to FIBAT (ROI→*I*) in (Figure 1b) to obtain a set of separated ROIs {*ROIs*|*τ*\*}. If any outputted set {*ROIs|τ*\*} contains more than one separated ROIs, then each ROI in the set {*ROIs*| *τ*\*} is further separated using the same procedure, thus forming a recursive loop. Otherwise, if any outputted set {*ROIs*|*τ*\*} contains only one ROI, then the recursion terminates. The final output of the local thresholding step *{ROIs}*_*n*_ is the union of all such sets containing one ROI that cannot be further separated.

### Fluorescence Intensity Based Adaptive Thresholding (FIBAT)

As described, FIBAT (Figure 1b) is used in both the global and local adaptive thresholding steps of each iteration of ACSAT to identify potential ROIs in the time-collapsed image *I* = *I*_*n*_ or to separate potentially overlapping neurons within *I* which is an element of 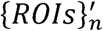, respectively. In either case, an optimal pixel intensity threshold value *τ*\* separates ROIs from the background. FIBAT selects *τ\** by searching for the threshold value that maximizes the number of resulting ROIs that are larger in area than *A*_*min*_ and smaller in area than *A*_*max*_. For the global adaptive thresholding step, we chose *A*_*min*_ = 50*px* ≈ 86*μm*^*2*^ and *A*_*max*_ = 300px ≈ 516*μm*^*2*^. For the local adaptive thresholding step, we chose *A*_*min*_ = 20*px* ≈ *μm*^*2*^ and *A*_*max*_ = ∞ because ROIs tend to shrink in size after repeatedly applying FIBAT.

The search is performed recursively over a pixel intensity range (*τ*_1_,*τ*_2_), where initially *τ*_*1*_ is the minimum pixel intensity value in *I* and *τ*_*T*_ is the maximum pixel intensity value in *I*. From this search range, *T* test threshold values *τ*_1_, *τ*_2_, …., *τ*_T−1_, *τ*_T_ are uniformly selected. A larger *T* will decrease the probability of skipping the optimal threshold value, but it will result in more computation time that may not be necessary. We chose *T* = 12. Each of these test threshold values *τ*_1_, …, *τ*_*T*_ is applied to the image *I* by assigning each pixel a 1 (a true calcium event) if its value is greater than the threshold or a 0 (a false calcium event) otherwise. Morphological operations are then performed to refine the thresholded images. Specifically, these operations fill in holes (0s surrounded by 1s) and remove spur pixels which may be due to noise. The operations also break H-connected ROIs prior to splitting overlapping cells. ROIs are finally collected with 8-connectivity (MATLAB function bwlabel or bwconncomp) to generate a set of segmented ROIs for each test threshold value: {*ROIs*|*τ*_1_}, …,{*ROIs|τ*_T_}.

Since ROIs represent real neurons that are roughly spherical in shape and are about 5 μm – 20μm in diameter, some realistic criteria can be used to eliminate false ROIs that are not possibly actual neurons. Accordingly, FIBAT removes ROIs from {*ROIs|**τ*_1_}, …, *{ROIs|τ*_*T*_} if their centroid is outside the ROI, or if their area is less than *A_min_* or greater than *A*_*max*_, or if their solidity (i.e. the area ratio between the convex hull of a ROI and the ROI itself) is greater than approximately the golden ratio.

The next search range is selected based on the results of the test thresholds. A relationship of the test threshold values *τ*_1_, …, *τ*_*T*_ vs the numbers of resulting ROIs |{*ROIs*|*τ*_1_}|, …, |{*ROIs*|*τ*_*T*_}| can be generated (Figure 1b). If the test threshold value *τ*_*c*_ resulted in the most ROIs i.e. *c* = arg max_c_ |{*ROIs|τ*_c_}|, then the next search range is set to (*τ*_max{1,c-1}_,*τ*_min{*T*,c+1}_) in order to include *τ*_*c*_ inside the search range. If more than one test threshold value *τ*_*c*_1__,*τ*_*c*_2__,… resulted in the same maximum number of ROIs, then the next search range is similarly set to (*τ*_max{1,min{*c*_1_,*c*_2_,….}−1}_, *τ*_min{*T*,max{*c*_1_,*c*_2_,….}+1}_) in order to contain all *τ*_*c*_1__, *τ*_*c*_2__, …. This search is terminated when further refinement of the search range produces little improvement in the number of detected ROIs: either the new search range |*τ*_*c*+1_ − *τ*_*c*−1_| is less than *ϵ* or the new range overlaps the previous range by at least *α*. We chose *α*= 90% and *ϵ* = the smallest non-zero intensity difference between every pair of adjacent pixels in whole image *I*. As such, *ϵ* is determined automatically and does not require user input. Upon termination, the optimal threshold value is set to 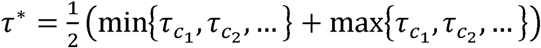, and the segmented ROIs {*ROIs*|*τ*\*} includes ROIs whose area exceeds *A*_*max*_.

### Code accessibility

The code/software described in the paper is freely available online at www.github.com/sshen8/acsat. The code is available as Extended Data.

## Results

We used the ACSAT algorithm (Figure 1a) to automatically segment ROIs from two datasets recorded with a custom wide-field microscope in the hippocampus and in the striatum of mice. The imaging area is 1.343×1.343 mm^2^, with a spatial resolution of 1024×1024 pixels (1.312×1.312μm^2^/pixel). Each dataset of approximately 4 GB was a continuous segment of a long image sequence acquired at 20 Hz with scientific CMOS camera for about 40 minutes. Each of the two datasets contained 2047 frames of 16 bits image corresponding to a recording duration of approximately 100 seconds.

Prior to the application of the ACSAT, the image sequences were time-collapsed as shown in Figure 2 and Figure 3 (top rows) for the hippocampal and the striatum datasets, respectively. These time-collapsed images show high intensity areas resembling neural morphology. The final segmented ROIs outputted by ACSAT are illustrated in Figure 2 and Figure 3 (bottom row), respectively. Time-collapsed images can be taken from longer sessions, but we found, in general, dynamic calcium events could be reasonably expected within this time window. A single calcium event from any given cell would yield a high intensity value in the time-collapsed images.

**Figure 2.**
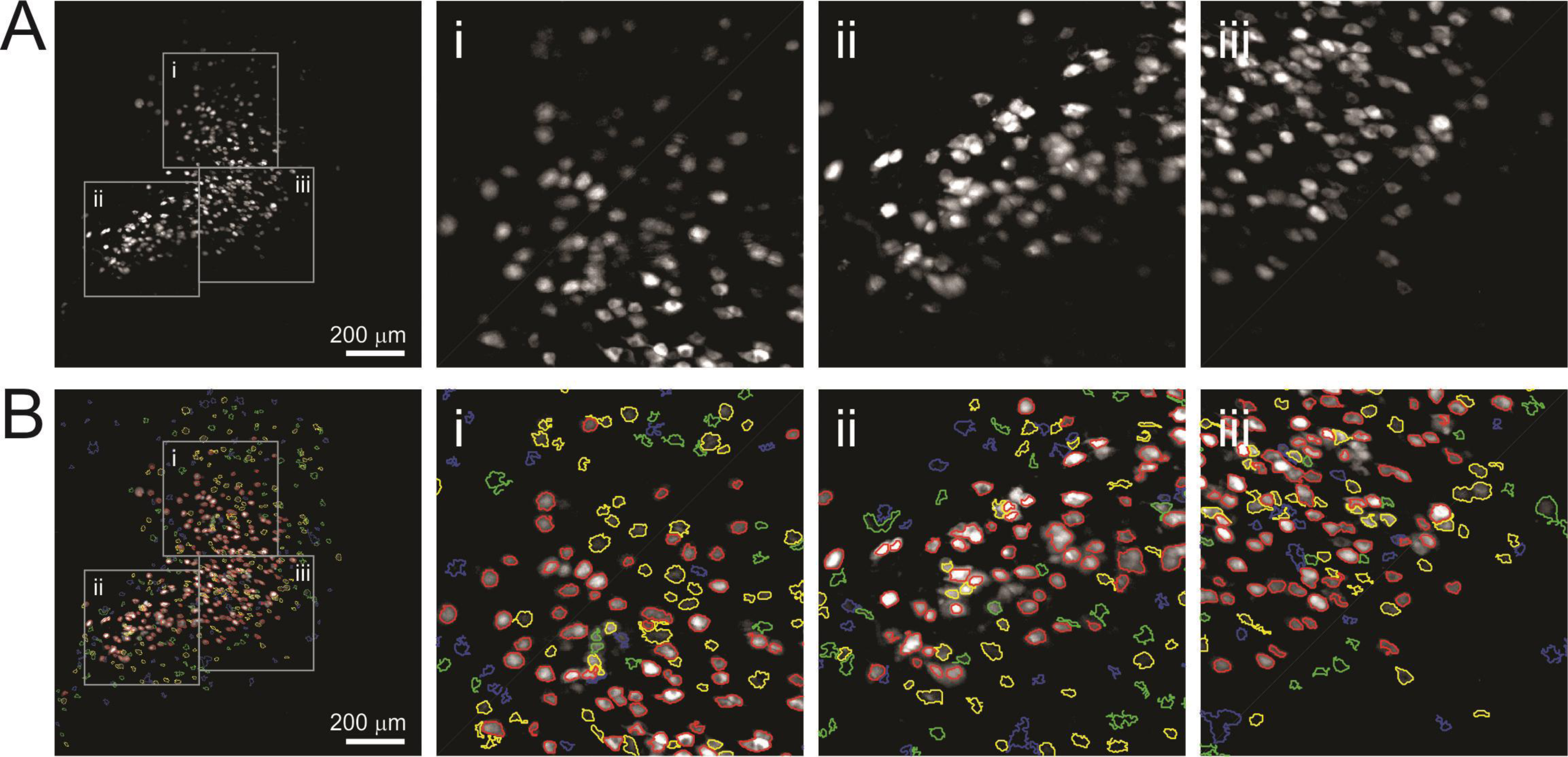
Hippocampus ROIs identified by ACSAT. (A) The aggregated image of hippocampus dataset and zoom-in detail images (Ai, Aii, and Aiii, corresponding to the grey boxes). (B) ACSAT ROIs from multiple iterations overlying on the aggregated image (red, yellow, green, and blue outline: iteration 1, 2, 3, and 4, respectively). The fourth iteration (blue) is shown for comparison although ACSAT terminated at iteration 3 (red, yellow, and green).

**Figure 3.**
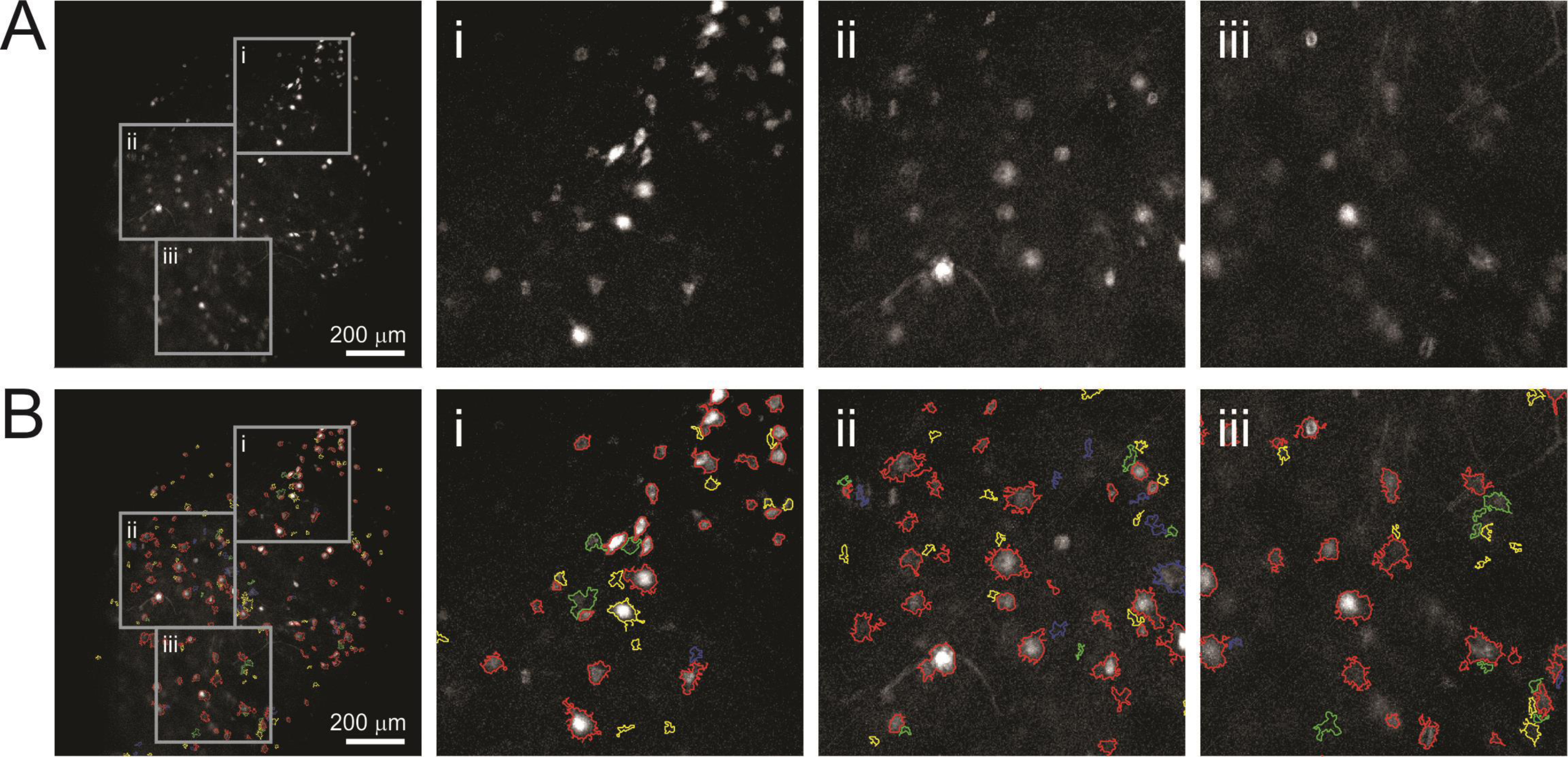
Striatum ROIs identified by ACSAT. (A) The aggregated image of striatum dataset and zoom-in detail images (Ai, Aii, and Aiii, corresponding to the grey boxes). (B) ACSAT ROIs from multiple iterations overlying on the aggregated image (red, yellow, green, and blue outline: iteration 1, 2, 3, and 4, respectively). The second (yellow), third (green), and fourth (blue) iterations are shown for comparison although ACSAT terminated at iteration 1 (red).

ACSAT is based on adaptive thresholding on a time-collapsed image, and thus it provides segmentation results at very fast speed. Specifically, to obtain the results as shown in Figure 2 and Figure 3, it took approximately one minute per iteration on a Xeon E5-1650 v4 at 3.6GHz with 128 GB DDR4 RAM, but it used less than 30 MB RAM. As such, the RAM size had little effect on the speed.

### ACSAT Accuracy Compared to Human-Generated Truth

To assess the accuracy of the ACSAT algorithm, we compared the ACSAT segmentation results with ROIs generated by human inspection (human-generated truth). Two human referees manually segmented ROIs from the raw image sequences. This human-generated truth contained 423 ROIs for the hippocampus dataset and 91 ROIs for the striatum dataset. We first compared the ACSAT-generated ROIs for the hippocampus and striatum datasets with the ROIs in the human-generated truth. We consider a pair of ROIs to correspond to the same neuron if they had centroids that were less than 50*px* ≈ 65.6*nm* apart and had a mutual overlap greater than 60%. We calculated the mutual overlap as the average of the percentages of the overlapping area against the areas of both ROIs. When there were multiple ROIs sharing overlapping areas, we selected the pair with highest mutual overlap as the matched ROIs.

For the hippocampus dataset, ACSAT identified 445 ROIs after three iterations. Among these 445 ROIs, 317 ROIs were matched in the human-generated truth (true positive), and 128 ROIs were not in the human-generated truth (false positive). Additionally, 106 ROIs in human-generated truth were not identified by ACSAT (false negative). This result gave us a precision rate of 71.2% (317 out of 445) and a recall rate of 74.9% (317 out of 423). For the striatum dataset, ACSAT was terminated after one iteration and identified a total of 135 ROIs: 69 true positives, 66 false positives, and 22 false negatives (precision rate: 51.1%, recall rate: 75.8%).

### ACSAT Accuracy beyond Human-Generated Truth

After a further examination of the individual ROIs identified by ACSAT that are false positives, our secondary manual inspection found that some of the false positive ROIs were actually true neurons that were missed in the initial human-generated truth due to human error. This means that ACSAT was able to segment ROIs that were not easily detectable by human experts. Specifically, for the hippocampus dataset, 70 (54.7%) out of 128 ROIs initially labeled as false positives were later determined to be actual neurons, and for the striatum dataset, 31 (47%) ROIs were true neurons. After correction, out of the total 445 ACSAT ROIs from the hippocampus dataset, 387 segmented ROIs corresponded to true neurons (true positive), and 58 segmented ROIs did not correspond to true neurons (false positive). Additionally, 106 true ROIs were not segmented (false negative). This corresponds to a precision rate of 87% and a recall rate of 78.5%. Similarly, for the striatum dataset, which resulted in 135 ACSAT ROIs, there were 100 true positives, 35 false positives, and 22 false negatives after correction. This corresponds to a precision rate of 74.1% and a recall rate of 82%. Even though neurons in the hippocampus and striatum have different morphology and dynamics, ACSAT was able to achieve the performance comparable to human referees for both datasets, and it was able to detect low-intensity neurons that were initially undetected by human referees. As such, our results demonstrate the robustness and effectiveness of the algorithm.

The result from the hippocampus dataset shows that ACSAT successfully identified true ROIs of diverse sizes (Figure 4, red). In general, the false positive ROIs had relatively smaller areas (Figure 4, yellow), similar to the ROIs missed by human referees (Figure 4, green). This indicates that ACSAT is more likely to recognize intensity changes in small areas thereby outperforming human referees under such challenging detection conditions. In the meantime, ACSAT missed a small portion of true ROIs, which shares similar sizes with those identified (Figure 4, blue).

**Figure 4.**
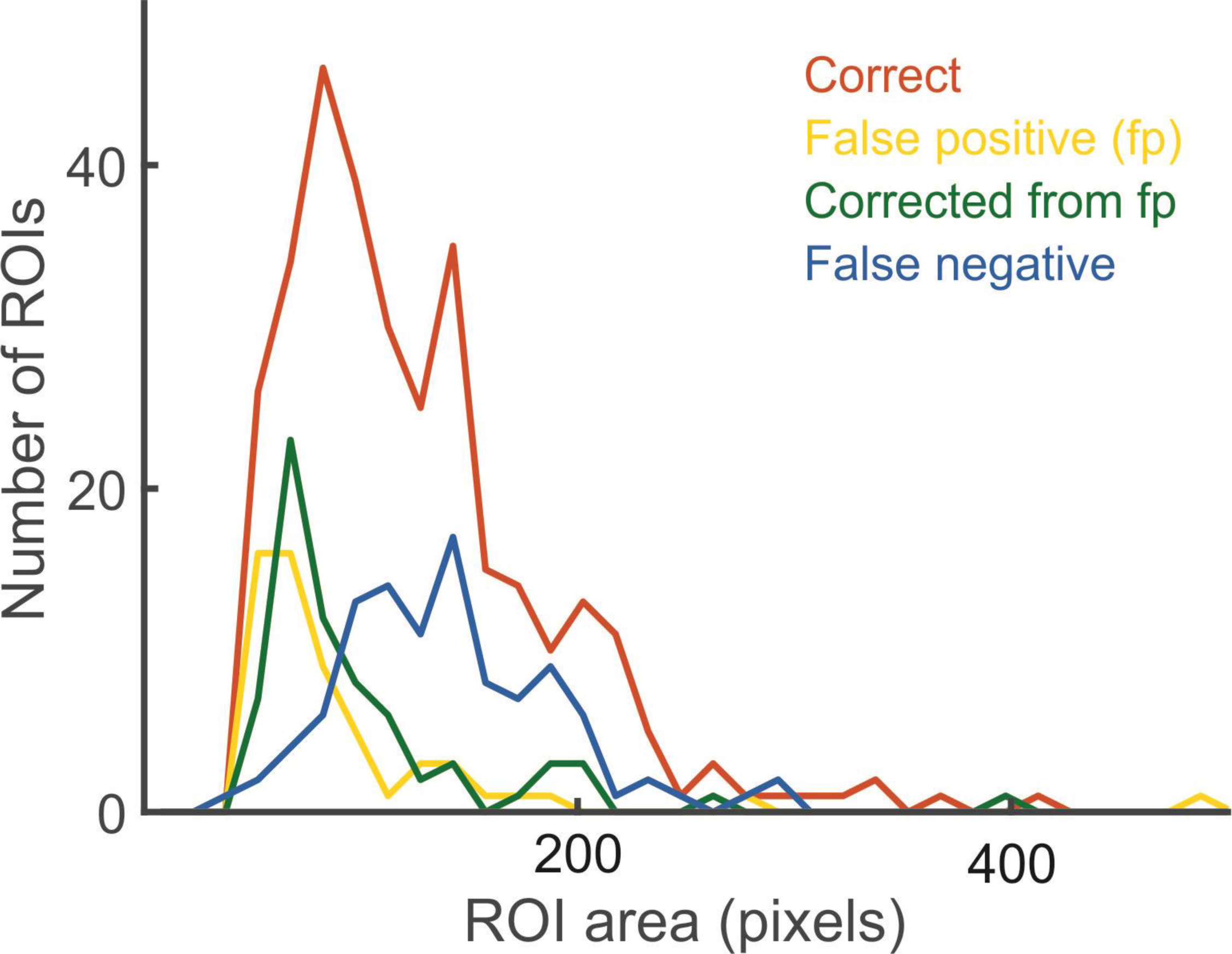
Distribution of ROI size. ACSAT identified true ROIs (red) with various size. The false positive ROIs (yellow) and those missed by human experts (green) tend to have small areas, while the areas of false negative ROIs (blue) leaned on the larger side.

### Number of Iterations in Using ACSAT

For the hippocampus dataset, ACSAT was terminated at iteration *n* = 3 when the change in global threshold value 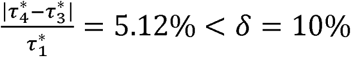. For the striatum dataset, ACSAT was terminated at iteration *n* = 1 when the change in global threshold value 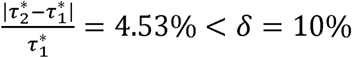.

To evaluate how ACSAT performs when terminated at different iteration numbers, we ran ACSAT up to 9 iterations on both datasets, and calculated several major performance indicators after each iteration (Figure 5): cumulative number of ROIs, global threshold value, recall, false negative rate, and false discovery rate (which is equal to 1 – precision) compared to the human-generated truth prior to secondary manual inspection of false positives. The cumulative number of ROIs, recall, and false discovery rate increased with the iteration number, but at different paces. While the cumulative number of ROIs and the false discovery rate increased steadily, recall rose steeply and reached its plateau within approximately three iterations for the hippocampus dataset and after the first iteration for the striatum dataset. Both the global threshold value and the false negative rate dropped as iterations progressed, indicating ACSAT dynamically adjusted the threshold to capture potential ROIs with lower intensity in later iterations. This dynamic adjustment of the threshold value was only possible because of the removal of segmented ROIs prior to each iteration. Overall, the changes in these performance indicators over iterations suggested that most true ROIs were identified during the early iterations: *n≤* 3 for the hippocampus dataset and *n* = 1 for the striatum dataset, which are consistent with when the ACSAT termination criterion described by *δ* was met. ROIs segmented during later iterations were mostly false positive.

**Figure 5.**
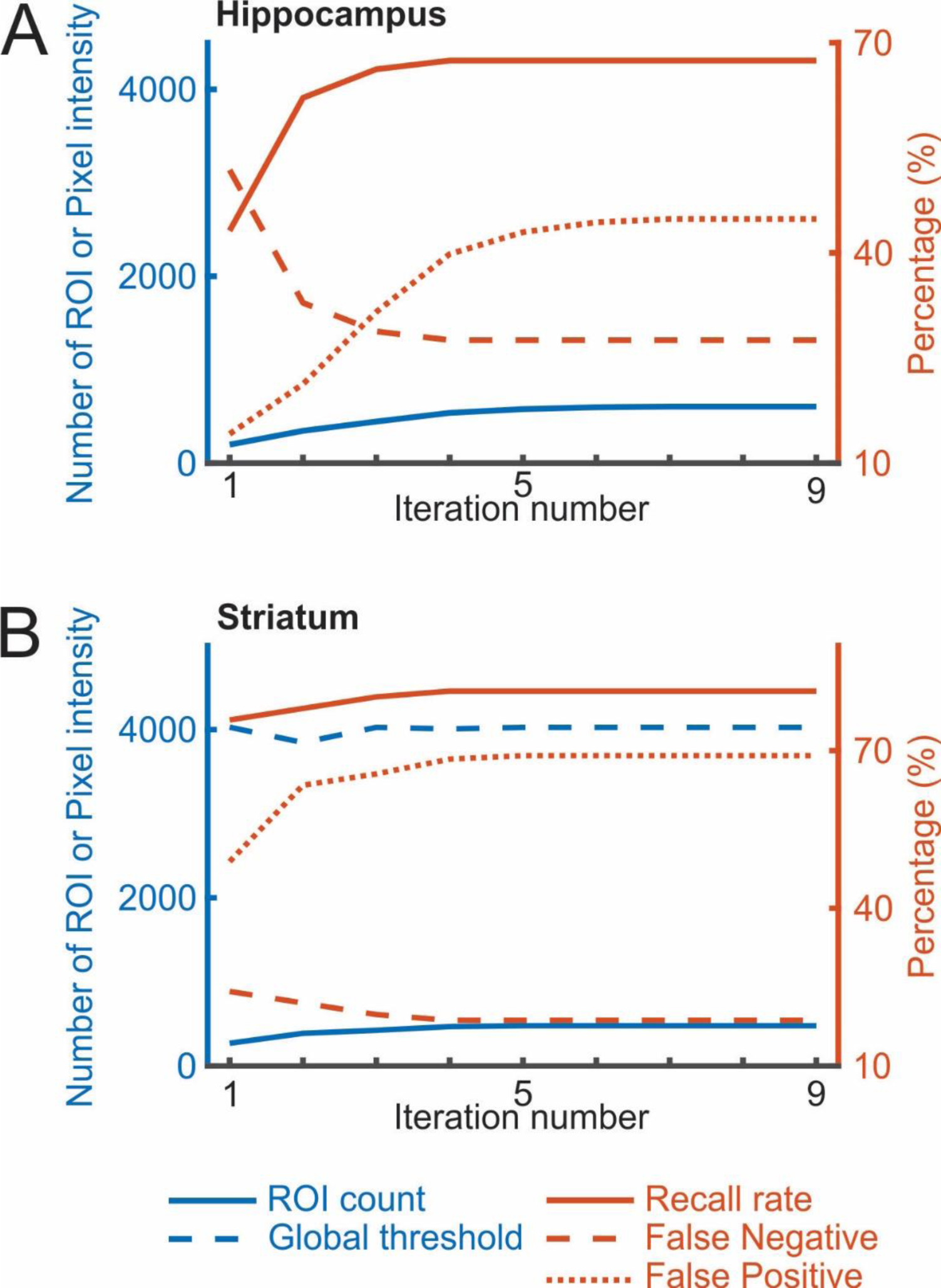
Cumulative performance of ACSAT over iterations. For both (A) hippocampus and (B) striatum datasets, the cumulative number of identified ROIs (blue line) increased steadily over iterations. The global threshold (dash blue line) tended to decrease with each iteration, allowing ACSAT to capture the ROIs with low intensity. Both recall (solid red line) and false discovery rate = 1 - precision (dotted red line) increased with iterations, while the false negative rate (dashed red line) decreased. All results reported here are based on human-generated truth prior to secondary manual inspection of false positives.

### FIBAT Global and Local Thresholding

In Figure 6, we demonstrate how FIBAT (Figure 1b) determines the threshold value that achieves optimal segmentation results by sampling the distribution of threshold value vs number of ROIs. Each trace of Figure 6 plots the number of ROIs that result from each sampled threshold value in the global thresholding step during the first four iterations of ACSAT (Figure 1a) on the hippocampus dataset. In each iteration, FIBAT (Figure 1b) first samples the threshold values across the entire intensity range at coarse resolution to identify the potential search range that may result in the maximum number of ROIs. FIBAT further re-samples threshold values within the new search range with a finer resolution, until it reaches a threshold value that gives the maximum number of ROIs. This design allows FIBAT to determine the optimal threshold value with a fine resolution without actually sampling the whole intensity range at the fine scale, and, as a result, reduces the processing time.

**Figure 6.**
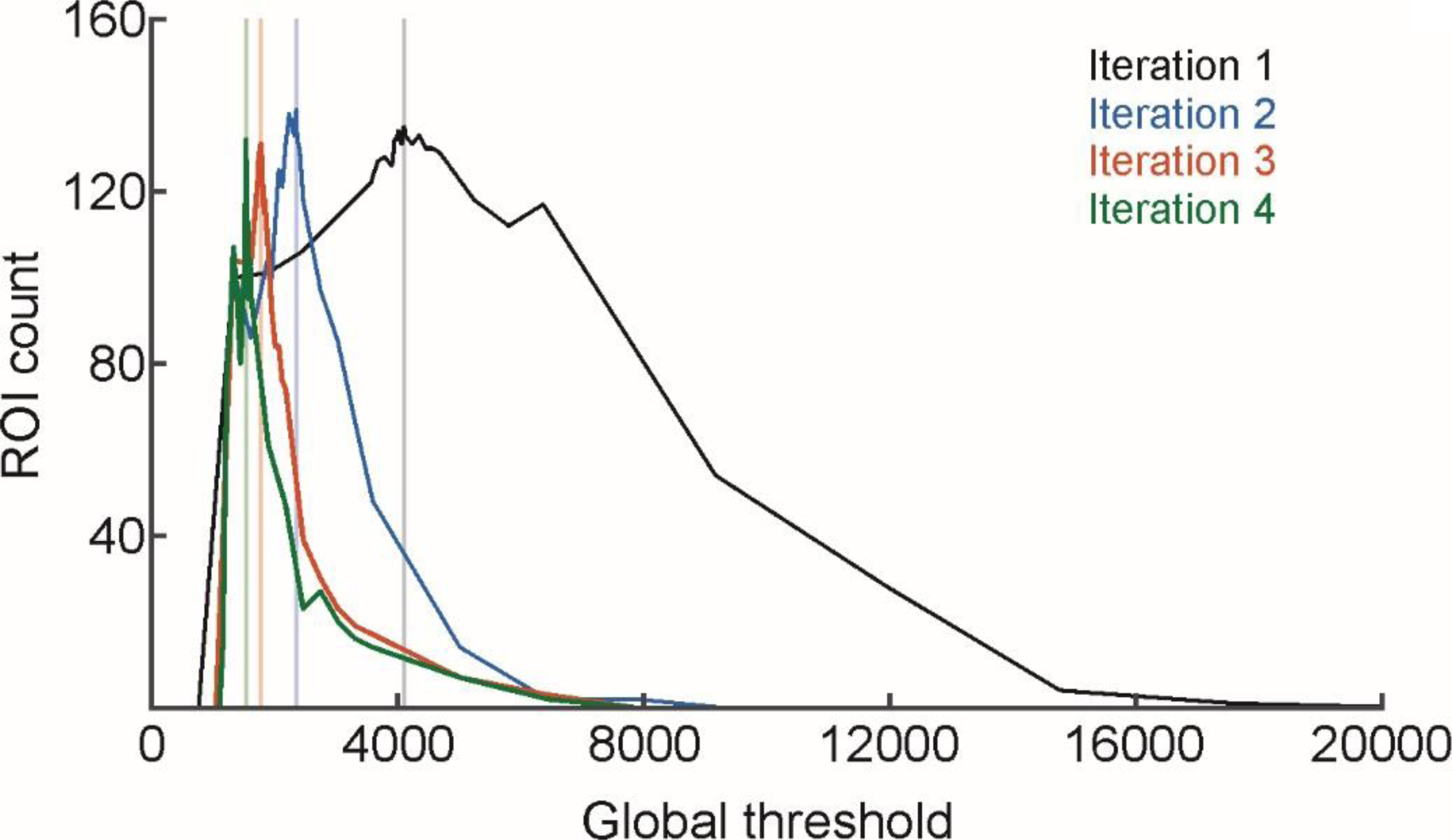
Convergence of the FIBAT optimal global threshold value for the hippocampus dataset. FIBAT first sampled at a coarse scale across a wide intensity range, and then focused on a small potential intensity range with a fine scale to identify the optimal global threshold value that generates most ROIs. The vertical lines indicate the final optimal global threshold value determined by FIBAT for each iteration.

After performing global thresholding to identify potential ROIs 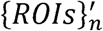 (Figure 1a), ACSAT further applies FIBAT locally to each identified ROI in 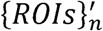 to refine the segmentation results (Figure 7). When neurons are densely labeled with GCaMP6, using the global thresholding step alone may lead to one or more large clusters of adjacent neurons being segmented as a single ROI (Figure 7a). For each such cluster, FIBAT (Figure 1b) determines and applies a new threshold value to the local ROI area. With local thresholding, the example cluster is further segmented into 5 ROIs (Figure 7b), which would not otherwise be separated by applying the global threshold. Because further local thresholding produces the same result (Figure 7c), the local thresholding step of ACSAT concludes that these 5 ROIs cannot be further separated, exits the recursive loop, and outputs these ROIs.

**Figure 7.**
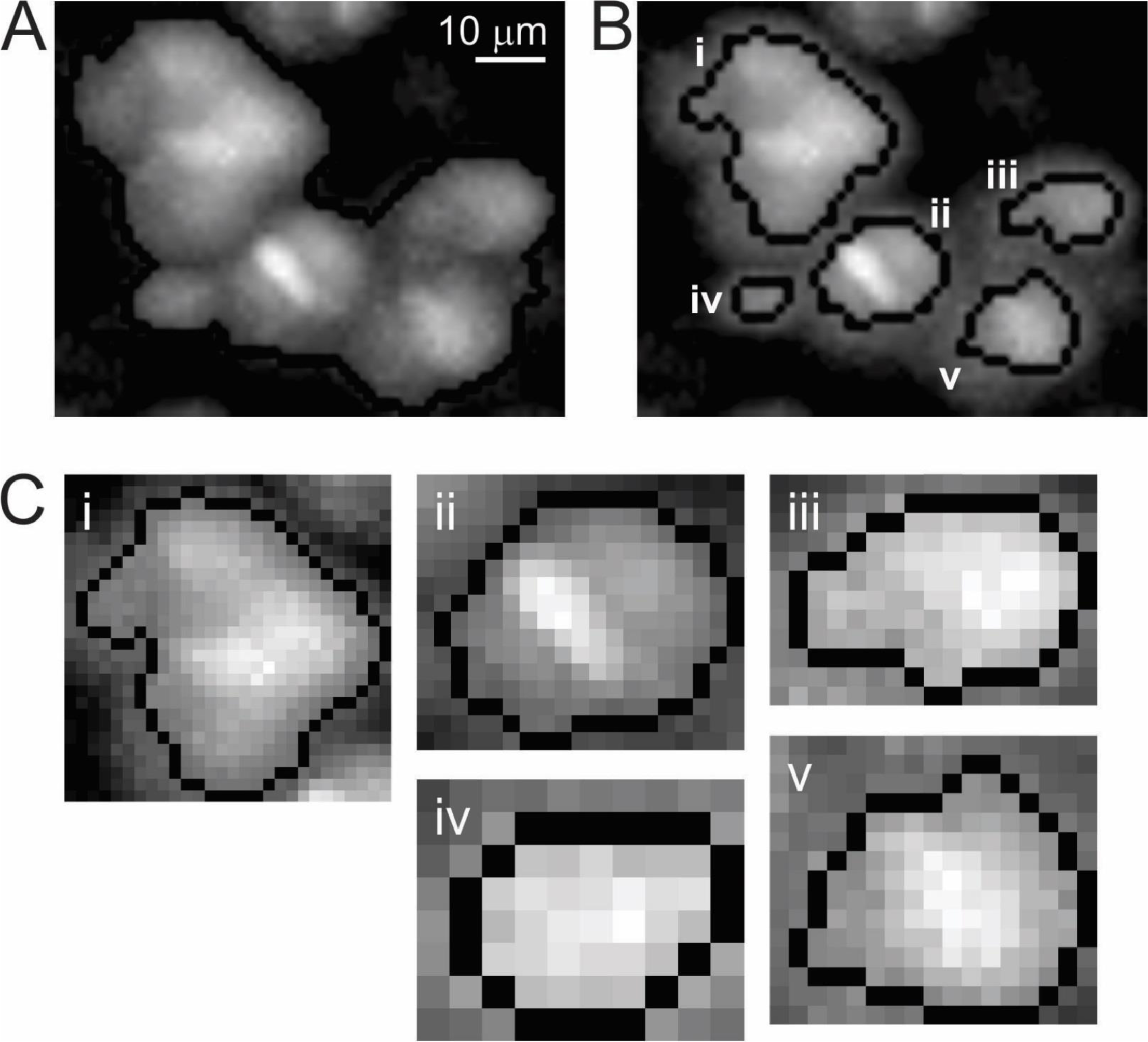
Improved ROI identification by local thresholding. (A) With global thresholding alone, a cluster of hippocampal neurons was identified as a single ROI. (B) After application of local thresholding, ACSAT successfully separated five ROIs from the single ROI. (C) Zoom-in of each ROI separated by local thresholding.

To evaluate the efficacy of local thresholding, we examined the hippocampus dataset at each iteration before and after the local thresholding step (Figure 8, left and right bars, respectively). Local thresholding refined the ROIs detected by global thresholding and captured more true ROIs at every iteration. It is also worth noting that, at later iterations, local thresholding was still able to identify true ROIs that were missed by global thresholding alone (Figure 8, iteration 4).

**Figure 8.**
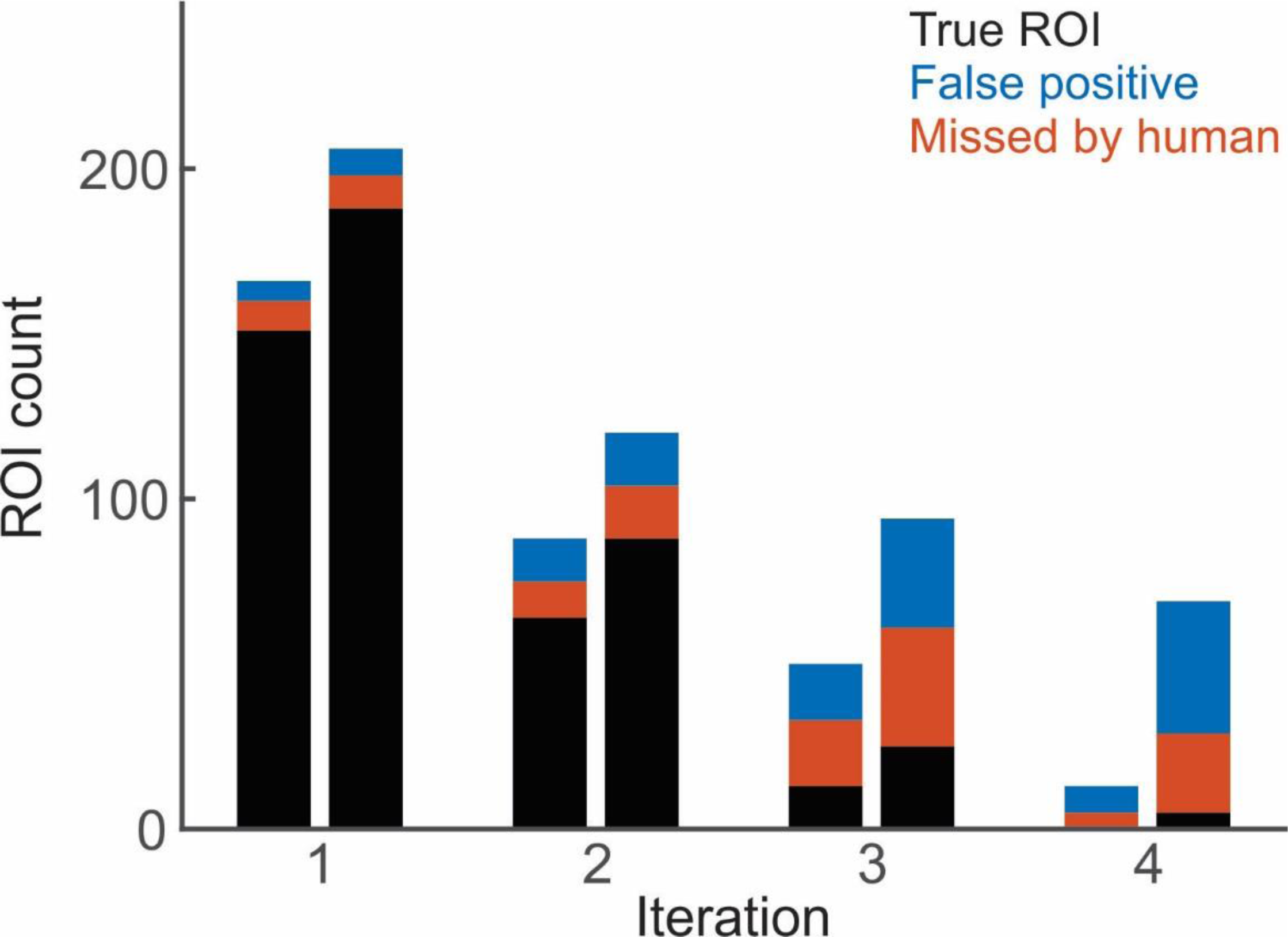
Local thresholding improves ACSAT performance. The ROIs identified by ACSAT at each iteration, before local thresholding (left) and after (right). Local thresholding separates overlapping ROIs and thus helped identify more ROIs, including those identified by human (black) or missed by human (red).

## Discussion

In this study, we presented our Automated Cell Segmentation by Adaptive Thresholding (ACSAT) method that adaptively selects threshold values based on image pixel intensity with two iterative steps at the global and local levels of time-collapsed image representing an image sequence. As such, the algorithm is capable of handling morphological and dynamic changes in neurons and is robust against luminance condition changes. When applied to two datasets collected from the hippocampus and the striatum in mice, ACSAT resulted in approximately 80% recall rate of regions of interest (ROIs) containing individual neurons (78.5% for the hippocampus dataset and 82% for the striatum dataset), and approximately 80% precision rate (87% for the hippocampus dataset and 74.1% for the striatum dataset). ACSAT was also able to detect low-intensity ROIs that were initially undetected by human referees.

The ACSAT algorithm is innovative because it is an intuitive thresholding method that uses global and local schemes to address variations in fluorescence intensity levels of GCaMP even within the same image field. Simply applying a lower global threshold value would result in few large ROIs containing multiple neurons within one ROI. On the other hand, with a high global threshold value, only a small number of neurons with high intensity would be found. As such, applying a single high or low threshold value would generate inadequate results with fewer or excessive ROIs, which is a universal limitation of thresholding methods. Our algorithm efficiently addresses this challenge in two ways. First, it cumulatively excludes previously segmented ROIs from the time-collapsed image *I*_*n*_ after each iteration so that in the following iteration, ACSAT could detect new ROIs that require distinct thresholds to separate but were missed with previous thresholds. Therefore, the global threshold value 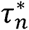 (Figure 1) used by ACSAT usually decreases after each iteration, and ROIs with high intensity were segmented before those with low intensity, as shown in Figure 2 and Figure 3. Because ACSAT is based on adaptive thresholding, it allows us to objectively and robustly segment ROIs with low intensity relative to the background. These low-intensity areas often pose challenges to human experts when manually detecting ROIs, as our results showed that about half of the ROIs initially labeled as false positive were actually true neurons (Figure 8).

Second, ACSAT uses FIBAT locally to separate overlapping ROIs one-by-one. This local approach directly addresses the issue that the intensities of pixels surrounding an ROI can vary. This heterogeneity of recorded neural signals is characteristic and common in biological data. However, because a higher thresholding value is usually required to separate adjoining neurons, the outputted sub-ROIs are often smaller than the corresponding true neurons. Because ROIs generally have pixel intensity decreasing radially, this shrinking effect is expected to be uniform around an ROI and thus can be corrected by a simple dilation.

Another major advantage of the ACSAT algorithm is its use of a time-collapsed image instead of the full image sequence during iterations. This makes our algorithm fast, less memory-demanding, and accurate as demonstrated by segmenting real recordings from mouse hippocampus and striatum. The global thresholding step generally took approximately 5 seconds to apply FIBAT to our full 1024 × 1024 pixels images *I*_*n*_. Because the local thresholding step uses FIBAT to separate sub-ROIs from ROIs that are usually much smaller than a full image, the local thresholding step is also relatively fast, taking approximately one minute per ACSAT iteration for the datasets reported. Neither of these two steps requires large memory because only the time-collapsed image is processed.

ACSAT has three sets of free parameters that can be objectively chosen or are not sensitive: *δ* which describes the termination condition for ACSAT, *α* which describes a termination condition for FIBAT, and *A*_*min*_ and *A*_*max*_ which describe the allowed sizes of ROIs. The latter set of parameters *A*_*min*_ and *A*_*max*_ are only determined by the experiment and data collection instrument, and should be chosen based on characteristics of the data acquisition conditions determined by, for example, image resolution and objective lens. The selection of other parameters should have minimal effect on ACSAT’s final segmentation results as discussed below.

The neuron ROI size criteria *A*_*min*_ and *A*_*max*_ should be chosen based on how large neurons are expected to be using information including image resolution, magnification, etc. The presence of these area criteria in our algorithm is consistent with the literature (Fantuzzo et al., 2017).

The termination condition for ACSAT, described by 5, can be explained by the tendencies of ACSAT. Specifically, running ACSAT for more iterations increases the number of ROIs segmented, especially the number of low-intensity ROIs, as the global threshold value 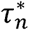 gradually decreases (Figure 5). While many of the added ROIs are true ROIs, the proportion of false positive ROIs added increases as iteration number increases (Figure 5). This increasing proportion of outputted false positives in later iterations can be attributed to the higher probability of a spurious collection of adjacent background pixels meeting the criteria to be an ROI. Also, the added false positives can be related to the step which clears previously segmented ROIs from the time-collapsed image at the start of each iteration of ACSAT. Due to the scattering of light in brain tissue, ROI removal may leave a few small fragments of brighter pixels around removed areas, which could be identified as ROIs during the next iteration. Most of the time, these misidentified ROIs were discarded either because of their small size or because they do not meet the solidity criteria; however, occasionally they may pass the size criteria and become the false positive ROIs. As a result, the majority of false positives tend to have small size (Figure 4, yellow).

In order to balance the effects of simultaneous increase in true ROIs and false positive ROIs, ACSAT stops when a decrease of global threshold value becomes relatively small between iterations i.e. 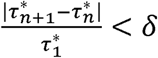. At that stage, most true ROIs have been detected and removed from the time-collapsed image. Thus, the global threshold values 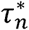 of any further iterations are similar, so most ROIs detected at this stage are false positives. For the hippocampus dataset, iteration *n =* 3 is when the increase in false positives begins to outweigh the increase in true positives, and for the striatum dataset, nearly all true ROIs segmented by ACSAT were outputted at iteration *n* = 1 (Figure 5). Qualitatively, the time-collapsed image *I*_0_ for hippocampus has a higher density of neurons with a greater variety of pixel intensities than the *I*_0_ for striatum, so it may take more iterations for ACSAT to perform at the same rate on the hippocampus dataset than on the striatum dataset. ACSAT’s performance under the diverse conditions of these two datasets suggests that our choice of *δ* = 10% provides a robust and objective termination condition for ACSAT that can be generalized to other datasets as well.

Additionally, the final segmentation results outputted by ACSAT are not very sensitive to the termination conditions for FIBAT described by *α* and *ϵ*. FIBAT is terminated if the threshold search range has minimal change over an iteration, which we determine in two ways. One way this condition would be satisfied is when all threshold values within the search range result in the same, optimal number of ROIs. This is equivalent to setting the criteria *α* = 100%. For the practical purpose of reducing FIBAT run time, we allow termination if the change in the search range is less than 1 – *α* = 10%. This condition is also trivially satisfied when FIBAT is used in the local thresholding step because, by definition, ROIs that cannot be separated by FIBAT will return exactly one ROI no matter what threshold value is used. Additionally, we terminate FIBAT if the search range is smaller than *ϵ*, the smallest difference between any pair of adjacent pixels in *I*, which can be objectively and automatically determined from *I*. If FIBAT were to continue refining the threshold value, then the gained precision beyond that defined by *ϵ* would be useless due to the discrete step in pixel intensity values in *I*.

The images *I*_0_ used by ACSAT are time-collapsed across whole time series, and therefore do not contain any temporal information. With the flexibility of ACSAT, the framework of ACSAT can be used with any other images that contain temporal information. A simple example is to define another input image 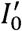 where the value of each pixel represents the time of its maximum intensity. This image 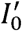 would allow ACSAT to separate adjoined ROIs that have similar intensity values in *I*_0_ but reach their maximum intensity at different time points, which is described by 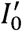. Other ways to generate the inputted image include correlations between nearby pixels, intensity dynamics such as standard deviation or variance, and/or combine them all. Overall, by taking advantage of adaptively determining the threshold value at both the global and local levels, ACSAT can theoretically perform segmentation on any image containing ROIs with non-homogenous intensity.

In this study, we evaluated the ACSAT’s performance with two GCaMP datasets; however, it is plausible to apply ACSAT on any imaging data, such as time-series data from voltage imaging, or even a regular static image, to identify neurons in the data.

Extended Data 1. ZIP file contains eleven MATLAB files.

Contributions: S.P.S. designed research, performed research, contributed unpublished reagents/analytic tools, analyzed data, and wrote the paper; H.T. performed research, analyzed data, and wrote the paper; K.R.H. performed research; R.W. analyzed data; H.G. performed research; J.S. and X.H. designed research and wrote the paper.

